# (p)ppGpp and CodY promote *Enterococcus faecalis* virulence in a murine model of catheter-associated urinary tract infection

**DOI:** 10.1101/655118

**Authors:** C Colomer-Winter, AL Flores-Mireles, S Kundra, SJ Hultgren, JA Lemos

## Abstract

In Firmicutes, the nutrient-sensing regulators (p)ppGpp, the effector molecule of the stringent response, and CodY work in tandem to maintain bacterial fitness during infection. Here, we tested (p)ppGpp and *codY* mutant strains of *Enterococcus faecalis* in a catheter-associated urinary tract infections (CAUTI) mouse model and used global transcriptional analysis to investigate the (p)ppGpp and CodY relationship. Absence of (p)ppGpp or single inactivation of *codY* led to lower bacterial loads in catheterized bladders, and diminished biofilm formation on fibrinogen-coated surfaces under *in vitro* and *in vivo* conditions. Single inactivation of the bifunctional (p)ppGpp synthetase/hydrolase *rel* did not affect virulence supporting previous evidence that association of (p)ppGpp with enterococcal virulence is not dependent on activation of the stringent response. Inactivation of *codY* in the (p)ppGpp^0^ strain restored *E. faecalis* virulence in the CAUTI model as well as the ability to form biofilms *in vitro*. Transcriptome analysis revealed that inactivation of *codY* restores, for the most part, the dysregulated metabolism of (p)ppGpp^0^ cells. While a clear linkage between (p)ppGpp and CodY with expression of virulence factors could not be established, targeted transcriptional analysis indicate that a possible association between (p)ppGpp and c-di-AMP signaling pathways in response to the conditions found in the bladder may plays a role in enterococcal CAUTI. Collectively, this study identifies the (p)ppGpp-CodY network as an important contributor to enterococcal virulence in catheterized mouse bladder and supports that basal (p)ppGpp pools and CodY promote virulence through maintenance of a balanced metabolism during adverse conditions.

**Importance:** Catheter-associated urinary tract infections (CAUTI) are one of the most frequent types of infection found in the hospital setting that can develop into serious and potentially fatal bloodstream infections. One of the infectious agents that frequently cause complicated CAUTI is the bacterium *Enterococcus faecalis*, a leading cause of hospital-acquired infections that are often difficult to treat due to the exceptional multidrug resistance of some isolates. Understanding the mechanisms by which *E. faecalis* causes CAUTI will aid in the discovery of new druggable targets to treat these infections. In this study, we report the importance of two nutrient-sensing bacterial regulators, named (p)ppGpp and CodY, for the ability of *E. faecalis* to infect the catheterized bladder of mice.

## Introduction

Catheter-associated urinary tract infections (CAUTI) are one of the most common hospital-acquired infections, accounting worldwide for about 40% of all nosocomial infections (1-3). In addition to substantially increasing hospitalization time and costs, CAUTI can lead to serious and potentially deadly secondary bloodstream infections (4). Complicated CAUTI is often the result of bacteria forming biofilms on indwelling urinary catheters, and enterococci (mainly *Enterococcus faecalis* and *E. faecium*) appear as the second leading cause of complicated CAUTI in many healthcare facilities (4-6). In addition, *E. faecalis* and *E. faecium* are major etiological agents of other life-threatening infections such as infective endocarditis, and an even more serious threat to public health due to their exceptional antibiotic resistance (7). The recent rise in enterococcal infections urges the development of new therapies, and understanding the mechanisms that promote *E. faecalis* CAUTI might uncover new druggable targets.

The pathogenic potential of *E. faecalis*, and of all enterococci in general, is tightly associated with their outstanding ability to survive an array of physical and chemical stresses, including common detergents and antiseptics, fluctuations in temperature, pH, humidity, and prolonged starvation (7). The regulatory second messengers ppGpp (guanosine tetraphosphate) and pppGpp (guanosine pentaphosphate), collectively known as (p)ppGpp, broadly promote bacterial stress tolerance and virulence (8, 9). In *E. faecalis*, (p)ppGpp has been shown to promote virulence in invertebrate and vertebrate animal models, and to mediate expression of virulence-related traits such as growth in blood and serum, biofilm formation, intraphagocytic survival and antibiotic tolerance (10-17). Originally described as the mediator of the stringent response (SR) (8), (p)ppGpp has distinct effects on bacterial physiology; at low (basal) concentrations it fine-tunes bacterial metabolism to adjust cellular growth in response to mild environmental changes (18), whereas at high levels it activates the SR responsible for promoting cell survival by slowing down growth-associated pathways and activating stress survival pathways (8, 18).

Two enzymes, the bifunctional synthetase/hydrolase Rel and the small alarmone synthetase RelQ, are responsible for enterococcal (p)ppGpp turnover (10, 19). Despite both Δ*rel* and Δ*rel*Δ*relQ* strains being unable to mount the SR (10, 13), there are fundamental differences in basal (p)ppGpp levels between these two strains. Specifically, while the double mutant, herein (p)ppGpp^0^ strain, is completely unable to synthesize (p)ppGpp, basal (p)ppGpp pools are about 4-fold higher in the Δ*rel* strain due to the constitutively and weak synthetase activity of RelQ (10, 14, 19). Accumulated evidence indicates that the metabolic control exerted by basal (p)ppGpp pools is more important during enterococcal infections than the semi-dormancy state characteristic of the SR (10, 13-17). This is exemplified by the distinct virulence phenotypes of Δ*rel* and (p)ppGpp^0^ strains, both unable to mount the SR . Specifically, while only the (p)ppGpp^0^ strain displayed attenuated virulence in *Caenorhabditis elegans* (10), *Galleria mellonella* (11, 13, 16) and in a rabbit abscess model (15), the Δ*rel* single mutant strain showed attenuated virulence in a rabbit model of infective endocarditis (17).

In low-GC Gram-positive bacteria such as *E. faecalis*, (p)ppGpp controls transcription of nutrient uptake and amino acid biosynthesis genes via the branched-chain amino acid (BCAA)- and GTP-sensing CodY regulator (20). It follows that (p)ppGpp accumulation during BCAA starvation severely depletes intracellular GTP pools in all Firmicutes such that CodY regulation is severely impaired due to depletion of its two co-factors (20). We recently confirmed the existence of the (p)ppGpp-CodY network in *E. faecalis* and demonstrated that inactivation of *codY* restored several phenotypes of the (p)ppGpp^0^ mutant strain, including virulence in *G. mellonella* (16). However, the contribution of the global nutritional regulator CodY to enterococcal pathogenesis in mammalian hosts remains unknown.

In this work, we examined the contribution of the (p)ppGpp and CodY to the pathogenesis of *E. faecalis* in a murine CAUTI model (21). We discovered that, in separate, basal levels of (p)ppGpp and the transcriptional regulator CodY promote biofilm formation in urine under *in vitro* conditions as well as virulence in a murine CAUTI model. Global transcriptome analysis validate earlier findings that deletion of *codY* restores, at least in part, the dysregulated metabolism of the cell in the absence of (p)ppGpp. Finally, targeted mRNA quantifications reveal that the (p)ppGpp/CodY network alters expression of genes coding for cyclic di-adenosine monophosphate (c-di-AMP) biosynthetic enzymes and of CAUTI virulence factors. Altogether, this study identifies the (p)ppGpp-CodY network as a contributor to enterococcal catheter colonization in the urinary tract and further supports that basal levels of (p)ppGpp promote bacterial virulence through maintenance of a balanced bacterial metabolism.

## Results

### The (p)ppGpp^0^ and Δ*codY* strains show impaired colonization in a murine CAUTI model

We compared the ability of the *E. faecalis* OG1RF parent, Δ*rel*, Δ*relQ*, (p)ppGpp^0^, Δ*codY* and (p)ppGpp^0^Δ*codY* strains to colonize and persist in the bladder of catheterized mice. Briefly, catheters were implanted into the bladder of C57BL/6Ncr mice prior to transurethral inoculation with ∼ 2 × 10^7^ CFU of each designated strain. Three days post-infection, the OG1RF strain was readily recovered from bladders (6.3 ± 0.6 log_10_ CFU) and catheters (6.0 ± 0.3 log_10_ CFU) of all infected animals (Fig. 1). Inactivation of *rel* (Δ*rel*) or *relQ* (Δ*relQ*) did not significantly alter *E. faecalis* colonization in bladders, but recovery of Δ*relQ* from catheters was significantly lower (∼ 0.8 log_10_ reduction, *p* =0.0027 when compared to OG1RF). On the other hand, the (p)ppGpp^0^ strain displayed ∼ 0.8 log_10_ (*p*=0.0015) and 1.5 log_10_ (*p*<0.0001) reductions in CFU recovered from bladders and catheters, respectively (Fig. 1). The Δ*codY* strain phenocopied the (p)ppGpp^0^ strain showing ∼ 0.8 log_10_ reductions in CFU recovered from both bladders (*p*=0.0156) and catheters (*p*=0.0031) (Fig. 1). However, inactivation of *codY* in the (p)ppGpp^0^ background [(p)ppGpp^0^Δ*codY* triple mutant] restored bacterial bladder and catheter colonization to near parent strain levels. In relative agreement with the bacterial loads detected on catheters and in bladders, only the OG1RF and Δ*rel* strains were consistently able to ascend and persist in mice kidneys three days post-infection (Fig. 1C).

**Figure 1.**
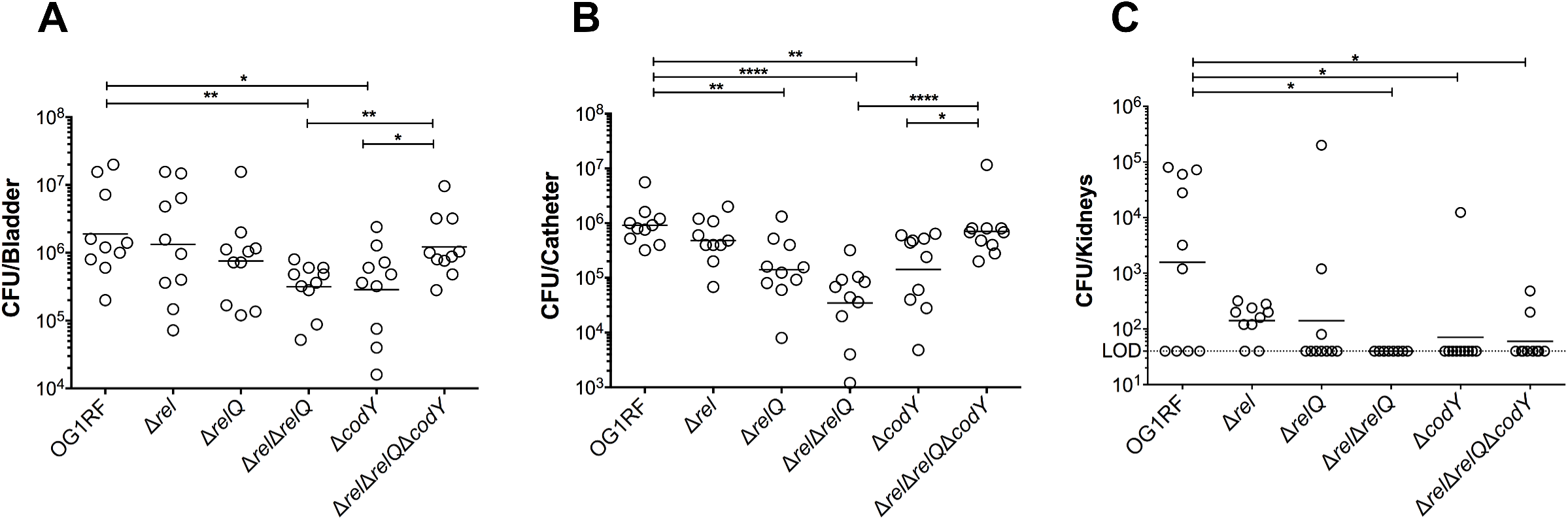
(p)ppGpp and CodY promote virulence of *E. faecalis* in a murine CAUTI model. The parent strain OG1RF and its derivatives were inoculated into the bladder of mice immediately after catheter implantation (n = 10). After 72 hours, animals were euthanized and bacterial burdens in (A) bladders, (B) catheters and (C) kidneys were quantified. Graphs show total CFU recovered from these sites, each symbol represents an individual mouse, and the median value is shown as a horizontal line. Symbols on the dashed line indicate that recovery was below the limit of detection (LOD, 40 CFU). The data was pooled from two independent experiments. Two-tailed Mann-Whitney *U* tests were performed to determine significance (\**p* < 0.05, \*\**p* < 0.005, \*\*\*\**p* < 0.0001).

### (p)ppGpp promotes timely growth of *E. faecalis* in human urine

The (p)ppGpp^0^ strain was previously shown to have growth and survival defects in whole blood and serum (15, 16). Interestingly, inactivation of *codY* in the (p)ppGpp^0^ background [(p)ppGpp^0^Δ*codY*] restored the (p)ppGpp^0^ growth defect in blood but not in serum (16). Here, we tested the ability of (p)ppGpp-defective (Δ*rel,* Δ*relQ* and (p)ppGpp^0^) and *codY* (Δ*codY* and (p)ppGpp^0^Δ*codY*) strains to replicate in pooled human urine *ex vivo*. Despite the colonization defect in the murine CAUTI model (Fig. 1), the Δ*codY* strain grew as well as the parent and Δ*relQ* strains (Fig. 2). On the other hand, Δ*rel*, (p)ppGpp^0^, and (p)ppGpp^0^Δ*codY* strains grew slower in the first 12 hours of incubation, with ∼ 0.6 log_10_ CFU difference at 3 and 7 hours post-inoculation (Fig. 2). Nevertheless, upon entering stationary phase, all strains reached similar growth yields and remained viable for at least 24 hours.

**Figure 2.**
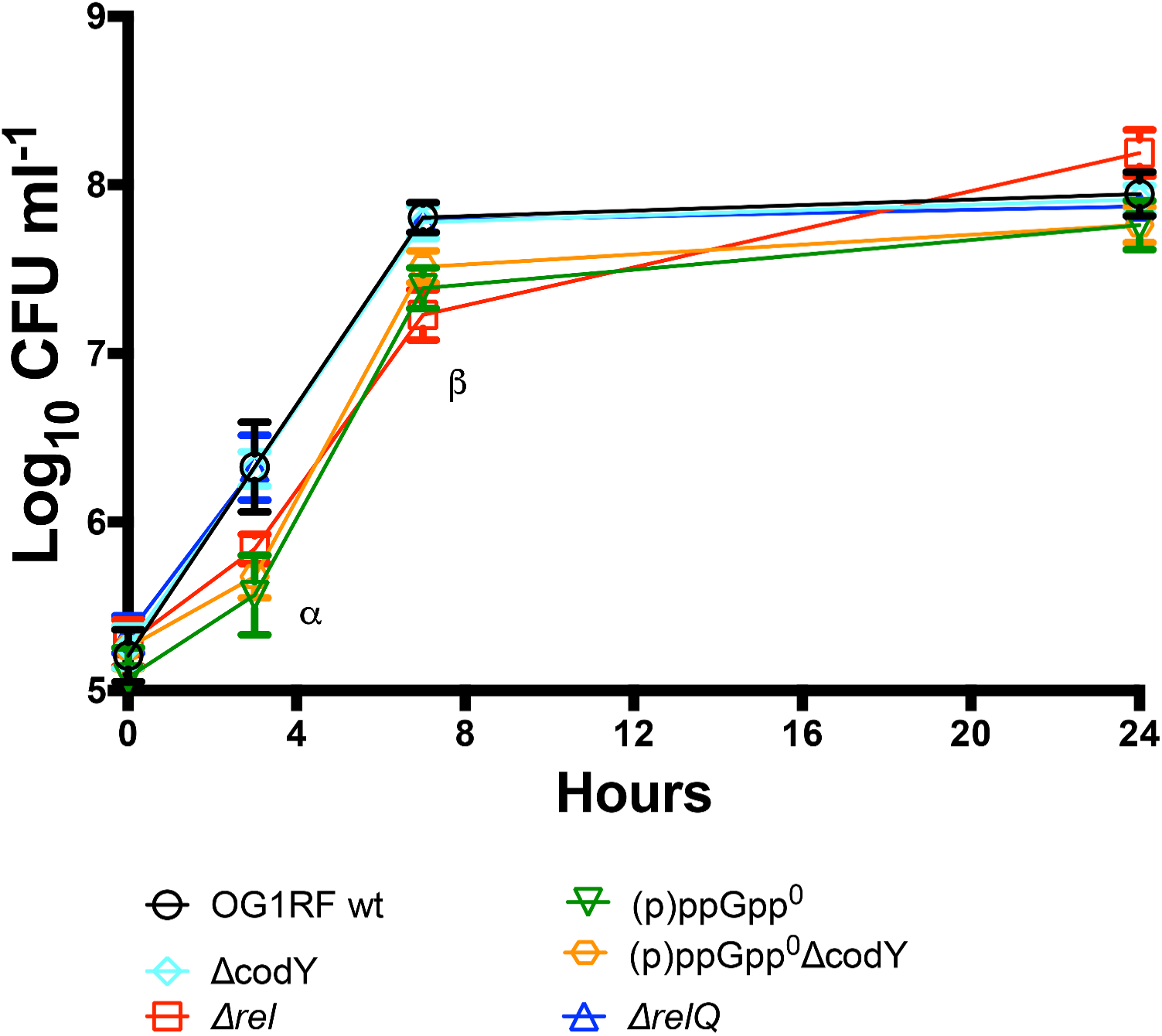
(p)ppGpp supports timely growth of *E. faecalis* in human urine. Growth of the parent *E. faecalis* OG1RF and its derivative mutant strains in pooled human urine. Aliquots at selected time points were serially diluted and plated on TSA plates for CFU enumeration. The graph shows the average log_10_-transformed CFU mean and standard deviations of three independent experiments. Mutant strains were compared to wild-type OG1RF by a two-way ANOVA with Dunnett’s multiple comparison test Asterisks indicate significant differences at 3 hours of incubation for Δ*rel,* (p)ppGpp^0^ and (p)ppGpp^0^Δ*codY* and at 7 hours of incubation for Δ*rel* and (p)ppGpp^0^Δ*codY* (*p* <0.0001).

### (p)ppGpp and CodY support biofilm formation in urine

Biofilm formation on urinary catheters is critical for enterococcal CAUTI (22-24). This is exemplified by the observation that *E. faecalis* OG1RF requires the presence of a catheter to persist for more than 48 hours in murine bladders (21). Follow up studies revealed that catheterization, in mice and in humans, triggers an inflammatory response that releases the host protein fibrinogen, which is used by *E. faecalis* as a nutrient as well as a scaffold to adhere and colonize the catheter surface (22-25). Taking into account that the reduction in bacterial counts of Δ*relQ*, (p)ppGpp^0^ and Δ*codY* strains was more pronounced on catheters than in bladders (Fig. 1) and that (p)ppGpp supports long-term survival of *E. faecalis* in biofilms (12), we sought to investigate if the attenuated virulence of these strains in murine CAUTI was linked to a reduced ability to form biofilms on fibrinogen-coated surfaces in the presence of urine. Since catheterization normally elicits proteinuria in the host, and *E. faecalis* requires a protein source to optimally form biofilms in urine (22, 24), human urine was supplemented with bovine serum albumin (BSA). In agreement with the bacterial loads obtained from the implanted catheters (Fig. 1B), quantifications of *E. faecalis* biofilm biomass revealed that Δ*relQ*, (p)ppGpp^0^ and Δ*codY* strains were deficient in biofilm production while the Δ*rel* strain phenocopied the parent OG1RF strain (Fig. 3A). Notably, inactivation of *codY* in the (p)ppGpp^0^ background alleviated the defective phenotype of the (p)ppGpp^0^ strain albeit the differences between the (p)ppGpp^0^Δ*codY* and OG1RF strains were still statistically significant.

**Figure 3.**
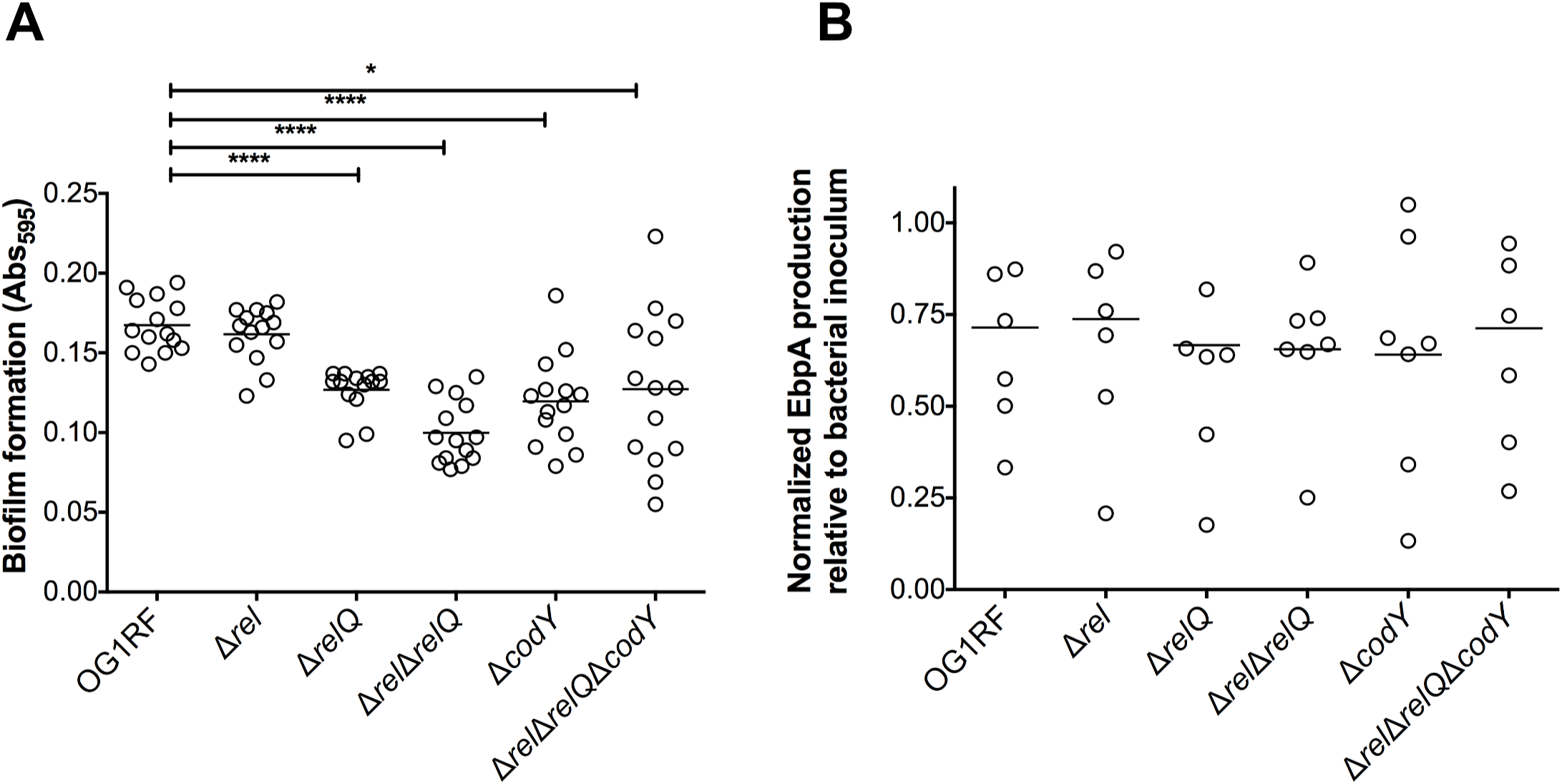
The (p)ppGpp/CodY network contributes to biofilm formation in urine. (A) Fibrinogen-coated 96-well polystyrene plates were incubated with *E. faecalis* strains for 48 hours in human urine supplemented with 20 mg ml^-1^ BSA. Plates were stained with 0.5% crystal violet, which was subsequently dissolved with 33% acetic acid and absorbance at 595 nm was measured. (B) EbpA quantification in urine. Strains were grown in urine + BSA overnight prior to quantification of EbpA by ELISA. EbpA surface exposure on *E. faecalis* cells was detected using mouse anti-EbpA^Full^ and HRP-conjugated goat anti-rabbit antisera, and absorbance was determined at 450 nm. EbpA production titers were normalized against the bacterial titers. Experiments were performed independently in triplicate and analyzed by a two-tailed Mann-Whitney *U* test (\**p* < 0.05, \*\*\*\**p* < 0.0001).

As indicated above, the main factor involved in *E. faecalis* biofilm formation on urinary catheters is production of the Ebp pilus, which binds directly to fibrinogen via the EbpA N-terminal domain (22, 26, 27). Quantification of EbpA via ELISA showed no noticeable reductions in EbpA production in any of the mutant strains (Fig. 3B) suggesting that the (p)ppGpp/CodY network promotes biofilm formation of fibrinogen-coated surfaces in an Ebp-independent manner.

### Inactivation of *codY* restores the dysregulated metabolism of the (p)ppGpp^0^ strain

Previously, we proposed that the virulence attenuation of the (p)ppGpp^0^ strain in different animal models is, in large part, due to the dysregulated metabolism caused by lack of (p)ppGpp control (13, 14, 16). By monitoring H_2_O_2_ production and the culture pH, we provided the first evidence that inactivation of *codY* restored a balanced metabolism to the (p)ppGpp^0^ background strain (16). To obtain additional insights into the (p)ppGpp-CodY relationship, we used RNA sequencing (RNA-seq) technology to compare the transcriptome of OG1RF, (p)ppGpp^0^ and (p)ppGpp^0^Δ*codY* strains grown to mid-exponential phase in the chemically-defined FMC medium (28) supplemented with 10 mM glucose (FMCG). In the (p)ppGpp^0^ strain, 690 genes were differentially expressed when compared to OG1RF (Table S1, *p* < 0.05, 2-fold cutoff), representing ∼ 27% of the entire *E. faecalis* OG1RF genome. In agreement with a previous microarray analysis (14), multiple PTS systems as well as citrate, glycerol, malate and serine utilization pathways were strongly induced in the (p)ppGpp^0^ strain under these conditions. A selected and representative number of dysregulated transport and metabolic genes in the (p)ppGpp^0^ strain are shown, respectively, in Tables 1 and 2. The upregulation of alternate carbon metabolism genes that are expected to be under carbon catabolite repression (CCR) in the glucose-rich condition of the FMCG medium indicates that complete lack of (p)ppGpp places *E. faecalis* in what we have originally termed a “transcriptionally-relaxed” state (14). When compared to the parent strain, 737 genes (∼ 29% of entire genome) were differentially expressed in the (p)ppGpp^0^Δ*codY* triple mutant strain (Table S2, *p* < 0.05, 2-fold cutoff). Despite the even large number of differentially expressed genes, inactivation of *codY* in the (p)ppGpp^0^ background strain normalized transcription of 274 genes that were dysregulated in the (p)ppGpp^0^ background (Table S3). The great majority (∼ 95%) of these genes were upregulated in the (p)ppGpp^0^ strain, supporting that deletion of *codY* abrogates, at least in part, the “transcriptionally-relaxed” state of the (p)ppGpp^0^ mutant. More specifically, inactivation of *codY* in the (p)ppGpp^0^ background normalized transcription (or brought to levels much closer to the parent strain) of ∼ 70 transport systems as well as over 130 metabolic genes including citrate, glycerol, malate, and serine utilization pathways, dehydrogenases, and molybdenum-dependent enzymes (Table S3).

**Table 1.** Transcriptional restoration of selected nutrient transport systems upon inactivation of *codY* in the (p)ppGpp^0^ background.

| Locus/ gene name | Function | Fold change*<br>(p)ppGpp<sup>0</sup> | Fold change<br>(p)ppGpp<sup>0</sup>Δ*codY* |
| --- | --- | --- | --- |
| PTS systems |  |  |  |
| OG1RF_10341 | PTS system | + 5.5 |  |
| OG1RF_10346 | PTS system | + 4.8 |  |
| OG1RF_10433 | PTS system | + 5.1 |  |
| OG1RF_10746 | PTS system | + 8.2 |  |
| OG1RF_11236 | PTS system | + 3.3 |  |
| OG1RF_11249 | PTS system | + 3.1 | - 3.3 |
| OG1RF_11512 | PTS system | + 5.0 |  |
| OG1RF_11614 | PTS system | + 4.2 |  |
| OG1RF_11780 | PTS system | + 3.8 |  |
| OG1RF_12261 | PTS system | + 4.6 |  |
| *bglP* | PTS system, β-glucoside uptake | + 7.4 |  |
| *frwB* | PTS system, fructose uptake | + 11.5 |  |
| *malX* | PTS system, maltose uptake | + 4.7 |  |
| *mltF2* | PTS system, mannitol uptake | + 7.1 |  |
| *scrA* | PTS system, β-glucoside uptake | + 3.4 |  |
| *sorA* | PTS system, sorbose uptake | + 32.5 |  |
| *treB* | PTS systems, trehalose uptake | + 11.9 | - 2.0 |
| *ulaB* | PTS system, ascorbate uptake | + 5.1 |  |
| ABC-type transporters |  |  |  |
| OG1RF_10665 | ABC-type transporter | + 3.4 | - 2.5 |
| OG1RF_10879 | ABC-type transporter | + 2.2 |  |
| OG1RF_11003 | ABC-type transporter | + 15.9 |  |
| OG1RF_11188 | ABC-type transporter | + 8.1 |  |
| OG1RF_11763 | ABC-type transporter | + 3.2 |  |
| OG1RF_11774 | Sugar ABC-type transporter | + 38.5 |  |
| OG1RF_12466 | ABC-type transporter | + 3.5 |  |
| *mdlB* | Multidrug ABC-type transporter | + 3.2 |  |
| *modB* | Molybdenum transporter | + 31.5 |  |
| T0 | Oligopeptide transporter | - 3.3 |  |
| *ziaA* | Zinc ABC-type transporter | + 3.2 |  |
| Other porters |  |  |  |
| OG1RF_11781 | Sodium symporter | + 5.3 |  |
| OG1RF_11873 | Phosphate transporter | + 4.5 | - 2.5 |
| OG1RF_11954 | Xanthine/Uracil permease | + 3.0 |  |
| OG1RF_12019 | Gluconate symporter | + 4.7 |  |
| OG1RF_12274 | Major Facilitator transporter | + 21.3 |  |
| OG1RF_12280 | Cytosine/purine permease | + 5.4 |  |
| OG1RF_12572 | Citrate transporter | + 51.8 |  |
| *dctM* | Organic Acid transporter | + 4.5 |  |
| *glpF2* | Glycerol uptake | + 11.0 |  |

(\*) Values represent fold-change in transcription when compared to the parent strain. All values shown were statistically significant ( $p < 0.05$ ). Blank fields indicate that there were no significant differences between mutant and parent strains.

**Table 2.** Transcriptional profile of selected metabolic genes upon inactivation of *codY* in the (p)ppGpp^0^ background.

| Locus/ gene name | Function | Fold change*<br>(p)ppGpp<sup>0</sup> | Fold change<br>(p)ppGpp<sup>0</sup>Δ*codY* |
| --- | --- | --- | --- |
| <u>Citrate metabolism</u> |  |  |  |
| OG1RF_10979 | Citrate carrier | + 31.2 |  |
| *citC* | Citrate metabolism | + 17.0 | - 5.0 |
| *citD* | Citrate metabolism | + 18.7 | - 3.3 |
| *citE* | Citrate metabolism | + 15.6 | - 5.0 |
| *citF* | Citrate metabolism | + 15.0 | - 5.0 |
| *citX* | Citrate metabolism | + 13.4 | - 5.0 |
| *citXG* | Citrate metabolism | + 13.2 | - 2.0 |
| <u>Serine metabolism</u> |  |  |  |
| *sdaA* | Serine dehydratase | + 103.5 |  |
| *sdaB* | Serine dehydratase | + 94.5 | - 2.5 |
| <u>Glycerol metabolism</u> |  |  |  |
| *glpK* | Glycerol kinase | + 5.7 | - 2.5 |
| *glpO* | Glycerol-3-P oxidase | + 6.6 | - 2.5 |
| *gldA2* | Glycerol dehydrogenase | + 13.7 |  |
| <u>Molybdenum metabolism</u> |  |  |  |
| OG1RF_11183 | MOSC protein | + 30.1 |  |
| OG1RF_11185 | Molybdopterin-binding protein | + 25.6 |  |
| OG1RF_11944 | Molybdenum hydroxylase | + 29.7 |  |
| OG1RF_11951 | Molybdenum hydroxylase | + 10.8 | - 2.0 |
| *moaA* | Molybdenum cofactor synthesis | + 20.0 |  |
| *moaB* | Molybdenum cofactor synthesis | + 27.1 |  |
| *moaC* | Molybdenum cofactor synthesis | + 10.1 |  |
| *yedF2* | Selenium metabolism | + 23.3 |  |
| *ygfJ* | Molybdenum hydroxylase | + 20.6 |  |
| <u>Other metabolic genes</u> |  |  |  |
| OG1RF_11942 | Ferredoxin NADP<sup>+</sup> reductase | + 23.4 |  |
| OG1RF_11943 | Flavodoxin | + 20.7 |  |
| *allD* | Ureidoglycolate dehydrogenase | + 14.7 |  |
| *mdh* | Malate dehydrogenase | + 16.5 |  |
| *fbp* | Fructose-1,6-bisphosphatase | + 10.6 | + 3.9 |
| *oadA* | Oxaloacetate decarboxylase | + 14.2 | - 3.3 |
| OG1RF_10107 | Glycosyl hydrolase | + 60.1 |  |
| *gcdB* | Glutaconyl-CoA decarboxylase | + 15.0 | - 5.0 |
| *arcC2* | Carbamate kinase | + 17.4 |  |
| *gltA* | Glutamate synthase | + 16.8 |  |
| OG1RF_11949 | Cysteine desulfurase | + 25.7 | - 2.5 |
| *hydA* | dihydropyrimidinase | + 14.6 |  |
| OG1RF_11585 | Ribosylpyrimidine nucleosidase | + 11.1 | - 3.3 |
| OG1RF_10108 | Endoribonuclease | + 48.7 |  |
| *endOF3* | N-acetylglucosaminidase | + 60.1 |  |

(\*) Values represent fold-change in transcription when compared to the parent strain. All values shown were statistically significant ( $p < 0.05$ ). Blank fields indicate that there were no significant differences between mutant and parent strains.

### Dysregulation of c-di-AMP homeostasis may affect fitness of (p)ppGpp^0^ and Δ*codY* strains in urine

Previously, the Ebp pilus and two high-affinity manganese transporters, EfaCBA and MntH2, were shown to be essential for *E. faecalis* virulence in a mouse CAUTI model (27, 29). In an attempt to identify (p)ppGpp- and CodY-dependent processes directly relevant to enterococcal CAUTI, we compared the transcriptional profile of these virulence factors in parent and mutant strains. In brief, quantitative real-time PCR (qPCR) was used to compare expression of selected genes in exponentially-grown BHI cultures of OG1RF, Δ*codY*, (p)ppGpp^0^ and (p)ppGpp^0^Δ*codY* strains before and after switching to pooled human urine. We found that transcription of *ebpA* (coding for the pilin sub-unit), *efaC* (the ATP-binding subunit of the ABC-type transporter EfaCBA) and *mntH2* was strongly induced upon shifting the parent strain culture from BHI to urine (Fig. 4). While full upregulation of *efaC* in urine appears to be dependent on CodY, *ebpA* and *mntH2* transcription was differentially impacted by (p)ppGpp and CodY. Specifically, while inactivation of CodY (Δ*codY*) limited induction of these genes after transition to urine, loss of (p)ppGpp induced transcription of *ebpA* and *mntH2* by ∼ 10-fold. Simultaneous inactivation of *codY* in the (p)ppGpp^0^ background [(p)ppGpp^0^Δ*codY*] modestly raised *ebpA* but fully restored *mntH2* mRNA levels (Fig. 4).

**Figure 4.**
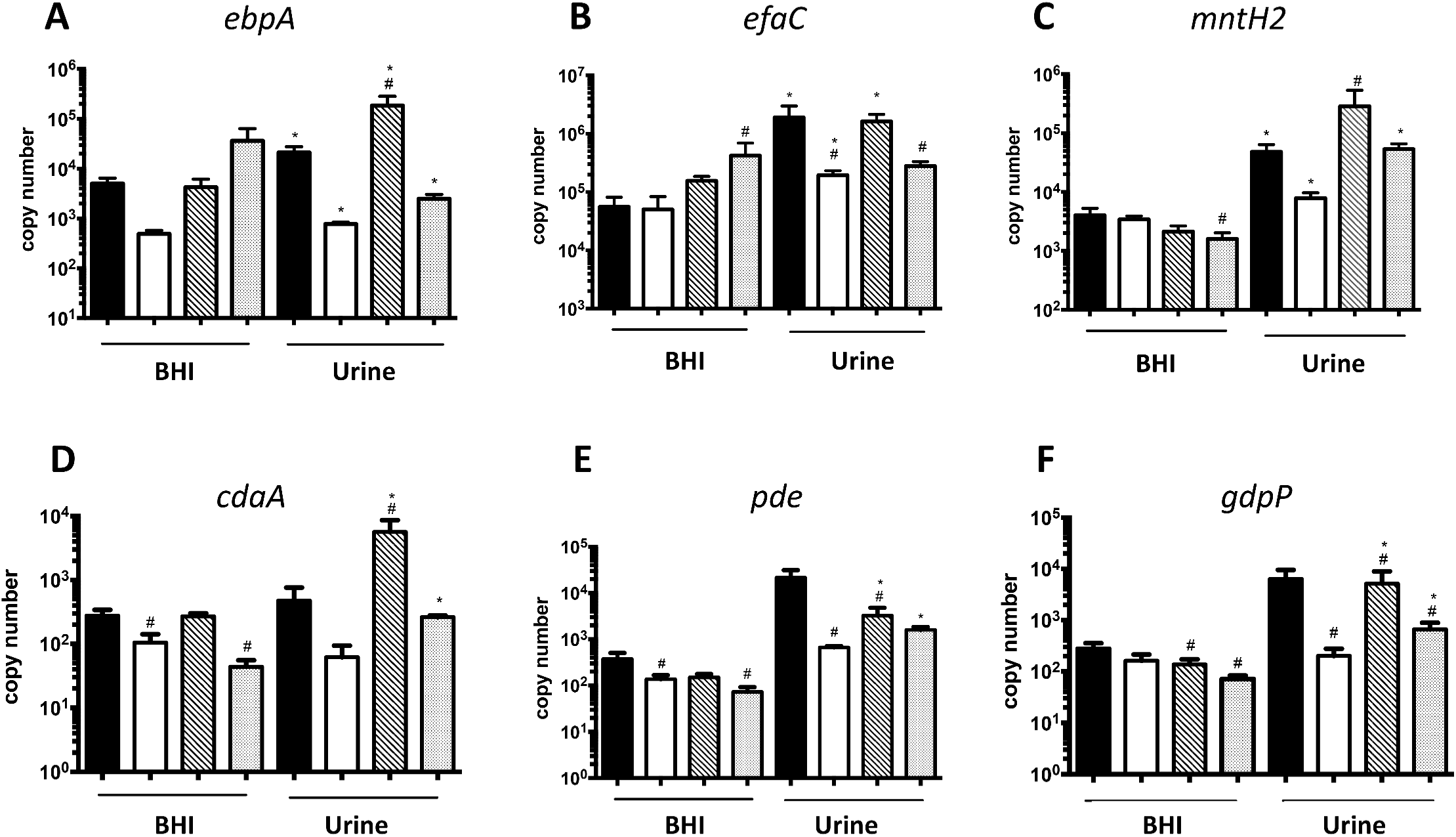
The transcript levels of *ebpA, efa, mntH2* and c-di-AMP synthase (*cdaA*) and hydrolases *(pde* and *gdpP)* genes in BHI and urine. The *E. faecalis* OG1RF wild type, (p)ppGpp^0^, (p)ppGpp^0^*ΔcodY* and *ΔcodY* strains were grown in BHI to mid-exponetial growth phase. Cell pellets were washed thoroughly and then exposed to pooled human urine and fresh BHI for 30 min. The transcript levels of (A) *ebpA*, (B) *efaC* (C) *mntH2* (D) *cdaA*, (E) *pde* and (F) *gdpP* were determined by quantitative real time-PCR (qRT-PCR). The bar graphs show average and standard deviations of results from three independent experiments performed in triplicate. Differences seen wuth the same strains under different conditions(#) or between parent and mutant strains under same growth condition (*) were compared via Student’s *t* test or ANOVA with multiple comparison test respectively, (p<0.05)

We also assessed transcription levels of genes coding for enzymes involved in c-di-AMP metabolism, a nucleotide second messenger essential for osmotic stress survival and whose regulatory network appears to be intertwined with the (p)ppGpp regulatory network in other bacteria (30-34). In the OG1RF strain, transcription of the c-di-AMP cyclase *cdaA* was not significantly altered upon transition from BHI to urine; however, transcription of both c-di-AMP hydrolases (*pde* and *gdpP*) was induced by ∼ 50- to 100-fold, respectively (Fig. 4). While induction of *gdpP* was dependent on CodY, activation of *pde* in urine was not as robust in all mutants strains. Unexpectedly, *cdaA* transcription was strongly induced (∼ 50-fold) in the (p)ppGpp^0^ strain when shifted from BHI to urine, suggesting that c-di-AMP levels may be dysregulated in the (p)ppGpp^0^ strain during CAUTI. Notably, c-di-AMP pools must be tightly controlled given that it is essential for growth but also highly toxic when present at high concentrations (35).

## Discussion

In this report, we showed that basal (p)ppGpp pools and the transcriptional regulator CodY mediate virulence of *E. faecalis* in a murine CAUTI model. These results corroborates previous findings that changes in basal levels of (p)ppGpp contribute to the virulence of *E. faecalis* (10, 11, 13, 15, 17) and show, for the first time, that CodY regulation is also important for the virulence of *E. faecalis*. Despite the attenuated virulence of the Δ*codY*, Δ*relQ* and (p)ppGpp^0^ strains, simultaneous inactivation of both regulatory pathways [(p)ppGpp^0^Δ*codY* strain] restored virulence to near parent strain levels. While this finding may appear contradictory, it is in line with previous studies showing that inactivation of *codY* restores virulence of (p)ppGpp^0^ strains in *Listeria monocytogenes* and *Staphylococcus aureus* (36, 37).

The association between (p)ppGpp and CodY, by which (p)ppGpp upregulates gene transcription via GTP depletion and concomitant alleviation of CodY repression, has been well established (20). We previously found that the (p)ppGpp and CodY regulatory networks are also intertwined in *E. faecalis* by showing that inactivation of *codY* restored phenotypes of the (p)ppGpp^0^ strain, such as the inability to grow in whole blood *ex vivo* and reduced virulence in the *G. mellonella* invertebrate model (16). By comparing final culture pH and H_2_O_2_ production of (p)ppGpp^0^ and (p)ppGpp^0^Δ*codY* strains, we hypothesized that deletion of *codY* restores a balanced metabolism to the (p)ppGpp^0^ strain (14, 16). The RNA-seq analysis reported here clearly supports this observation as multiple genes coding for alternate carbon utilization pathways were strongly upregulated in the (p)ppGpp^0^ strain but not in the (p)ppGpp^0^Δ*codY* triple mutant strain.

Our results also indicate that restoration of global metabolism in the (p)ppGpp^0^Δ*codY* strain when compared to the (p)ppGpp^0^ strain may be particularly relevant in experimental CAUTI. Specifically, we show that while absence of either (p)ppGpp or CodY results in defects in biofilm formation *in vitro* and *in vivo*, simultaneous inactivation of the (p)ppGpp and CodY regulatory systems restores CAUTI virulence and partially restores *in vitro* biofilm formation to near parent strain levels (Fig 1. and Fig. 3). Considering that the levels of the surface protein EbpA was not altered in any of the mutant strains (Fig. 3), we propose that (p)ppGpp and CodY promote biofilm formation on urinary catheters by mediating the metabolic rearrangements required for biofilm growth and survival in urine. In fact, a comparative transcriptome analysis reveals that several metabolic pathways previously identified as relevant to enterococcal adaptation to growth in urine (38) overlap with the (p)ppGpp and CodY regulons (13, 14, 20). For example, transcriptional activation of alternate carbon utilization and amino acid biosynthesis/transport genes in response to the relatively low urinary concentrations of amino acids and glucose (38-41) are shown here to be regulated by (p)ppGpp and CodY (Table S1, Table 1 and Table 2) (13, 14, 20).

In an attempt to identify specific (p)ppGpp- and CodY-regulated processes that are important for CAUTI, we used qPCR to compare transcription of known virulence factors such as the Ebp pilus and the metal transporters EfaCBA and MntH2 (27, 29) in parent and mutant strains. Although there were significant alterations in transcription of *ebpA, efaC* and *mntH2* in the (p)ppGpp and *codY* mutant strains, it was not possible to establish a firm correlation between the transcriptional expression of these virulence factors and the attenuated virulence phenotypes shown in Figure 1. Due to the association of c-di-AMP signaling with biofilm formation (33, 34) and adaptation to osmotic stress (31, 32) – two relevant traits during CAUTI – and the previous linkage with (p)ppGpp signaling (30, 35, 42), we similarly assessed gene expression of the enzymes responsible for c-di-AMP synthesis (*cdaA*) and degradation (*pde* and *gdpP*). c-di-AMP is an emerging regulatory nucleotide shown to control a number of cellular processes, including potassium homeostasis, osmotic adaptation and biofilm formation (35). In addition, previous investigations revealed an intricate but poorly understood association between the c-di-AMP and (p)ppGpp signaling networks, in which (p)ppGpp regulates c-di-AMP levels by serving as an allosteric regulator of c-di-AMP phosphodiesterase (PDE) enzymes, while c-di-AMP stimulates (p)ppGpp synthesis via an unknown mechanism (30, 35, 42). The strong activation of *pde* and *gdpP* in the parent strain upon shift to urine suggests that intracellular c-di-AMP levels decrease in the presence of urine. This is not surprising considering that salt concentrations in urine are generally high (average osmolality between 300 to 900 mOsm/kg H_2_O), and that low c-di-AMP pools are associated with bacterial salt tolerance (43) whereas increased c-di-AMP levels are linked to salt hypersensitivity (44-46). Interestingly, transcription of the c-di-AMP synthetase gene (*cdaA*) in urine was approximately 100-fold higher in the (p)ppGpp^0^ strain when compared to the parent strain, indicating that c-di-AMP homeostasis may be severely disrupted in the (p)ppGpp^0^ strain. Finally, transcription of the *pde* and *gdpP* genes was not as strongly induced (or not induced at all) in the (p)ppGpp^0^ and Δ*codY* strains. While the cellular levels c-di-AMP in cells grown in urine remain to be determined, and the possibility of the (p)ppGpp/c-di-AMP signaling crosstalk in *E. faecalis* confirmed, the qPCR analysis suggests that *E. faecalis* lowers c-di-AMP pools to adjust its metabolism to the environment (i.e. high osmolality) encountered in urine.

By comparing the genes restored by *codY* inactivation in our RNA-Seq analysis (Table S1, Table 1 and Table 2) to the transcriptome of *E. faecalis* OG1RF grown in urine (38), several other common pathways were identified providing additional leads as to why alleviation of CodY repression restored the biofilm defect of the (p)ppGpp^0^ strain. Specifically, Vebø *et al*. showed that transcription of a major glycerol uptake system was induced when the OG1RF strain was grown in urine *ex vivo*, suggesting that *E. faecalis* utilizes glycerol as a source of energy to grow and survive in urine (38). Notably, aerobic metabolism of glycerol by the GlpO enzyme is also the main source of H_2_O_2_ generation in *E. faecalis* (47) and *E. faecalis* activates transcription of several oxidative genes (such as *sodA, npr* and *trxB*) when grown in urine (38), possibly to cope with increased ROS generation caused by aerobic glycerol metabolism. In agreement with previous microarray data (14), the current RNA-seq analysis revealed that transcription of glycerol catabolic genes, such as *glpO and gldA2*, was activated by 6-fold or more in the (p)ppGpp^0^ strain (Table S1, Table 2), likely augmenting ROS production in the urinary tract. Another possibility, which is not mutually exclusive from the others, is that the strong (∼ 20-fold) upregulation of molybdenum metabolism genes in the (p)ppGpp^0^ strain is disadvantageous to *E. faecalis* when grown in the urinary tract environment. Molybdenum is a rare transition metal that functions as a co-factor of several redox active enzymes (48-50); it should be noted that urothione, the degradation product of the molybdenum cofactor in humans, is excreted in urine (51). In contrast to other metal co-factors, molybdenum is catalytically active only when complexed with a pterin-based scaffold, forming the Moco prosthetic group (50). The Moco biosynthetic genes that are highly induced in the (p)ppGpp^0^ mutant strain code for enzymes that require GTP and the oxygen-reactive iron and copper metals (50), possibly linking molybdenum metabolism to (p)ppGpp and oxidative stress. Finally, *E. faecalis* appears to downregulate serine catabolism in urine (38). However, serine dehydratases were among the most highly expressed genes in the (p)ppGpp^0^ strain (∼ 100-fold), likely favoring pyruvate generation instead of protein synthesis. While it is unknown how serine catabolism and dysregulation of other metabolic and transport pathways affect biofilm formation and virulence in *E. faecalis*, alleviation of CodY repression for the most part restored transcription of these genes to wild-type levels.

Collectively, the results presented here reveal that (p)ppGpp and CodY support *E. faecalis* presence in the catheterized murine urinary tract by controlling the metabolic arrangements necessary for the fitness of *E. faecalis* in this environment and, possibly, by modulating biofilm formation. Future studies using global transcriptome and metabolome analysis of (p)ppGpp-deficient and CodY-deficient strains recovered directly from CAUTI, coupled with characterization of pathways relevant to biofilm formation in urine, are necessary to fully understand how alleviation of CodY repression restores biofilm formation in the absence of (p)ppGpp. In addition, the relevance of c-di-AMP signaling to enterococcal CAUTI and the relationship between (p)ppGpp and c-di-AMP signaling pathways will warrant further investigations. In this regard, work is underway to determine the scope and targets of c-di-AMP regulation in *E. faecalis*. These studies should expand our mechanistic understanding of how global metabolic regulators such as CodY, (p)ppGpp and possibly c-di-AMP mediate enterococcal pathogenesis in the urinary tract and beyond.

## Materials and Methods

### Bacterial strains and growth conditions

The parent *E. faecalis* OG1RF strain and its derivatives Δ*rel,* Δ*relQ,* Δ*rel*Δ*relQ* [(p)ppGpp^0^], Δ*codY* and (p)ppGpp^0^Δ*codY* strains have been previously described (10, 16). All strains were routinely grown in BHI at 37°C. For RNA-seq analysis, overnight cultures of *E. faecalis* OG1RF, (p)ppGpp^0^, and (p)ppGpp^0^Δ*codY* strains were diluted in a 1:100 ratio in 50 ml of the chemically-defined FMC medium (28) supplemented with 10 mM glucose (FMCG) and allowed to grow statically at 37°C to an OD_600_ of 0.3. Growth in pooled human urine from healthy donors (Lee Biosolutions) was monitored as described elsewhere, with minor modifications (29). Briefly, overnight cultures were diluted 1:100 in PBS and inoculated 1:100 for growth assessment in urine. Cultures were incubated aerobically at 37°C and, at selected time-points, aliquots were serially diluted and plated on TSA plates for colony-forming unit (CFU) determination. To determine the transcriptional responses of selected genes upon transition from laboratory media to human urine, overnight cultures grown in BHI were diluted 1:20 in 5 ml of fresh sterile BHI and allowed to grow statically at 37°C to an OD_600_ of 0.5. The bacterial cells were washed twice with PBS (pH 7.0) and pelleted down by centrifuging at 2500 rpm for 8 min. After washing, pellets were resuspended in 7.5 ml filter sterilized urine and incubated at 37°C for 30 min. The controls were resuspended in the same volume of fresh BHI and incubated at 37°C for 30 min.

### Mouse catheter implantation and infection

The mice used in this study were 6-week-old female wild-type C57BL/6Ncr mice purchased from Charles River Laboratories. Mice were subjected to transurethral implantation and inoculated as previously described (21). Mice were anesthetized by inhalation of isoflurane and implanted with a 5-mm length platinum-cured silicone catheter. When indicated, mice were infected immediately following catheter implantation with 50 µl of ∼2 × 10^7^ CFU of bacteria in PBS introduced into the bladder lumen by transurethral inoculation as previously described (21). To harvest the catheters and organs, mice were sacrificed 3 days post-infection by cervical dislocation after anesthesia inhalation; silicone catheter, bladder and kidneys were aseptically harvested. The Washington University Animal Studies Committee approved all mouse infections and procedures as part of protocol number 20150226. All animal care was consistent with the Guide for the Care and Use of Laboratory Animals from the National Research Council and the USDA Animal Care Resource Guide.

### Biofilm formation in human urine

For assessment of biofilm formation on fibrinogen-coated 96-well polystyrene plates (Grenier CellSTAR), wells were coated overnight at 4°C with 100 µg ml^-1^ human fibrinogen free of plasminogen and von Willebrand Factor (Enzyme Research Laboratory). The next day, *E. faecalis* overnight cultures were diluted to an optical density (OD_600_) of 0.2 in BHI broth. The diluted cultures were centrifuged, washed three times with 1x PBS, and diluted 1:100 in urine supplemented with 20 mg ml^-1^ BSA. Bacterial cells were allowed to attach to the fibrinogen-coated plate at 37°C under static conditions as described in(29, 52). After 24 hours, microplates were washed with PBS to remove unbound bacteria and biofilm formation assessed by staining wells with crystal violet as previously described (52). Excess dye was removed by rinsing with sterile water and then plates were allowed to dry at room temperature. Biofilms were resuspended with 200 µl of 33% acetic acid and the absorbance at 595 nm was measured on a microplate reader (Molecular Devices). Experiments were performed independently in triplicate per condition and per experiment.

### Presence of EbpA on the cell surface of (p)ppGpp- and CodY-deficient strains

Surface expression of EbpA by *E. faecalis* OG1RF and derivatives was determined by ELISA as previously described (23). Bacterial strains were grown for 18 hours in urine supplemented with 20 mg ml^-1^ of BSA. Then bacterial cells were washed (3 times) with PBS, normalized to an OD_600_ of 0.5, resuspended with 50 mM carbonate buffer (pH 9.6) containing 0.1% sodium azide and used (100 µl) to coat Immulon 4HBX microtiter plates overnight at 4°C. The next day, plates were 3 times washed with PBS containing 0.05% Tween 20 (PBS-T) to remove unbound bacteria and blocked for 2 h with 1.5% bovine serum albumin (BSA)–0.1% sodium azide–PBS followed by three PBS-T washes. EbpA surface expression was detected using mouse anti-EbpA^Full^ antisera, which was diluted 1:100 in dilution buffer (PBS with 0.05% Tween 20, 0.1% BSA, and 0.5% methyl α-d-mannopyronoside) before serial dilutions were performed. A 100-μl volume was added to the plate, and the reaction mixture was incubated for 2 h. Subsequently, plates were washed with PBS-T, incubated for 1h with HRP-conjugated goat anti-rabbit antisera (1:2,000), and washed again with PBS-T. Detection was performed using a TMB substrate reagent set (BD). The reaction mixtures were incubated for 5 min to allow color to develop, and the reactions were then stopped by the addition of 1.0 M sulfuric acid. The absorbance was determined at 450 nm. Titers were defined by the last dilution with an A_450_ of at least 0.2. As an additional control, rabbit anti-*Streptococcus* group D antiserum was used to verify that whole cells of all strains were bound to the microtiter plates at similar levels. EbpA expression titers were normalized against the bacterial titers at the same dilution.

### RNA-seq analysis

Cells grown in FMCG to OD_600_ 0.3 were collected by centrifugation at 4000 rpm for 20 min at 4°C, and resuspended in 4 ml of sterile RNA stabilizing solution [3.5 M (NH_4_)_2_SO_4_, 16.6 mM sodium citrate, 13.3 mM EDTA, adjusted to a pH of 5.2 with H_2_SO_4_]. After a 10 min incubation at room temperature, cells were centrifuged at 4000 rpm for 30 min at 4°C, and pellets were stored at - 80°C. RNA was extracted using the hot acid-phenol:chloroform method previously described (53). Subsequently, precipitated RNA was treated once with DNase I (Ambion, Carlsbad, CA), followed by a second DNase I treatment using the DNA-free kit (Ambion) to completely remove DNA, divalent cations and traces of the DNase I enzyme (204). RNA concentrations were quantified with the NanoDrop 1000 spectrophotometer (NanoDrop, Wilmington, DE) and RNA quality assessed with the Agilent Bioanalyzer (Agilent, Santa Clara, CA). RNA sequencing (RNA-Seq), data processing and statistical analysis was performed at the University of Rochester Genomics Research Center (UR-GRC) using the Illumina platform as previously described (54). Gene expression data have been deposited in the NCBI Gene Expression Omnibus (GEO) database (www.ncbi.nlm.nih.gov/geo) under GEO Series Accession number GSE131749

### Real-time quantitative PCR analysis

Cultures were pelleted at 2500 rpm for 8 min at 4°C. The bacterial pellets were resuspended in 1 ml of RNA protect and incubated for 5 min at room temperature followed by centrifugation at 2500 rpm for 8 min at 4°C. At that point, the bacterial pellets were kept at -80°C until ready for RNA extraction. Cells were resuspended in TE buffer (10 mM Tris Cl [pH 8], 1mM EDTA) and 10 % SDS, homogenized for three 30 second cycles, with 2 min on ice between cycles. The nucleic acids were retrieved from the total protein by phenol:chloroform (5:1) extraction. The inorganic phase was resuspended in 0.7 ml RLT buffer (Qiagen) supplemented with 1 % β-mercaptoethanol, and RNA was purified using RNeasy mini kit (Qiagen), including the on-column DNase treatment recommended by the supplier. To further reduce the DNA contamination, RNA samples were treated with DNase I (Ambion) at 37°C for 30 min and were re-purified using RNeasy mini kit (Qiagen). RNA concentrations were determined using Nanodrop. Reverse transcription and real-time PCR were carried out according to protocols described previously (53) using the primer sets indicated in **Table S4**.

### Statistical Analysis

Data sets were analyzed using GraphPad Prism 6.0 software unless otherwise noted. Log_10_-transformed CFU values from urine growth curves were analyzed via a two-way ANOVA followed by comparison post-test. For CAUTI experiments, biofilm assays and EbpA ELISA, a Two-tailed Mann-Whitney *U* test was performed. RNA-seq processing and the subsequent statistical analysis was performed at the University of Rochester Genomics Research Center (UR-GRC) as previously described (54). qPCR data was analyzed by Student’s *t*-test and ANOVA with multiple comparisons test, respectively.

## Acknowledgements

This research was supported by grants from the National Institute of Allergy and Infectious Disease AI135158 (JAL) and AI10874901 (SJH), and the National Institute of Diabetes and Digestive Kidney Diseases DK051406 (SJH) and P50-DK0645400 (SJH). CCW was supported by the American Heart Association GSA Predoctoral Fellowship 16PRE29860000. The funders had no role in study design, data collection and analysis, decision to publish, or preparation of the manuscript.

## Supporting information

**Table S1.** List of genes differentially expressed in the (p)ppGpp^0^ strain in comparison to the parent OG1RF strain (*p* ≤ 0.05, 2-fold cutoff).

**Table S2.** List of genes differentially expressed in the triple mutant Δ*codY*(p)ppGpp^0^ strain in comparison to the parent OG1RF strain (*p* ≤ 0.05, 2-fold cutoff).

**Table S3.** Side by side comparison of linear fold-change of selected genes in the (p)ppGpp^0^ strain and triple mutant Δ*codY*(p)ppGpp^0^ strain in comparison to the parent OG1RF strain.

**Table S4.** Primers used for qPCR.

